# Genetic risk for Alzheimer’s disease predicts hippocampal volume through the lifespan

**DOI:** 10.1101/711689

**Authors:** Kristine B Walhovd, Anders M. Fjell, Øystein Sørensen, Athanasia Monica Mowinckel, Céline Sonja Reinbold, Ane-Victoria Idland, Leiv Otto Watne, Andre Franke, Valerijia Dobricic, Fabian Kilpert, Lars Bertram, Yunpeng Wang

**Affiliations:** Center for Lifespan Changes in Brain and Cognition, Department of Psychology, University of Oslo, POB 1094 Blindern, 0317 Oslo, Norway; Division of Radiology and Nuclear Medicine, Oslo University Hospital, Rikshospitalet, POB 4950 Nydalen, 0424 Oslo; Oslo Delirium Research Group, Department of Geriatric Medicine, University of Oslo, Norway, POB 4950 Nydalen, 0424 Oslo; Institute of Basic Medical Sciences, University of Oslo, Norway, POB 4950 Nydalen, 0424 Oslo; Institute of Clinical Molecular Biology, Christian-Albrechts-University of Kiel, Kiel, Germany; Lübeck Interdisciplinary platform for Genome Analytics, University of Lübeck, Maria-Goeppert-Str. 1 (MFC1), 23562 Lübeck, Germany

**Keywords:** Lifespan, Hippocampus, Polygenic risk score, APOE, Alzheimer’s Disease, Development, Aging, MRI

## Abstract

**INTRODUCTION:** It is unknown whether genetic risk for Alzheimer’s disease (AD) represents a stable influence on the brain from early in life, or whether effects are age-dependent. It is critical to characterize the effects of genetic risk factors on the primary neural substrate of AD, the hippocampus, throughout life.

**METHODS:** Relations of polygenic risk score (PGS) for AD, including variants in Apolipoprotein E (*APOE*) with hippocampal volume and its change were assessed in a healthy longitudinal lifespan sample (n = 1181, 4-95 years), followed for up to 11 years with a total of 2690 MRI scans.

**RESULTS:** AD-PGS showed a significant negative effect on hippocampal volume. Offset effects of AD-PGS and *APOE* ε4 were present in hippocampal development, and interactions between age and genetic risk on volume change were not consistently observed. DISCUSSION: Endophenotypic manifestation of polygenic risk for AD may be seen across the lifespan in healthy persons.

**Highlights:**

- Genetic risk for AD affects the hippocampus throughout the lifespan
- *APOE* ε4 carriers have smaller hippocampi in development
- Different effects of genetic risk at different ages were not consistently observed
- Genetic factors increasing risk for AD impact healthy persons throughout life
- A broader population and age range are relevant targets for attempts to prevent AD

## 1. Background

Among the earliest behavioral signs of Alzheimer’s disease (AD) are deficits in memory and orientation, closely tied to dysfunction of the hippocampus and its neural circuits [1]. While variants at multiple genetic loci identified in case-control studies are associated with increased AD risk [2], very little is known about how these genetic variants affect individuals devoid of AD diagnosis at an endophenotypic level. While select studies have shown that the major genetic risk factor for AD, the apolipoprotein E (*APOE*) ε4 allele [3, 4], is associated with smaller hippocampal volumes at various ages also in healthy persons [5], longitudinal data are scarce and restricted to older adults [6]. Recently, a small study (n = 66) indicated a relationship of AD genetic risk factors in addition to *APOE* ε4 and hippocampal atrophy in healthy older adults [7]. To date, it remains unknown whether polygenic risk scores (PGS) calculated from established AD risk variants translate to differences in neural characteristics at different life stages in healthy persons.

The negative effect of common genetic polymorphisms on late-life disease, including cognitive decline and onset of AD, has often been interpreted in terms of age-specific mechanisms. For instance, effects of the *APOE* ε4 allele have been understood within the framework of antagonistic pleiotropy [8, 9]. This term was coined by Williams [10], postulating genes that have opposite effects on fitness at different ages or in different somatic environments. In this manner, it has been assumed that such genetic risk variants for AD either have no effect until older age is reached, or that while they confer a risk in aging, they might present some advantage in development [8, 9]. However, there is growing evidence pointing to the continuous influence of early life factors for later-life cognitive function and its neural foundations [11–14]. This makes it crucial to investigate the extent to which genetic factors established to influence late life neural and cognitive disease act in a temporally stable and dimensional manner at the level of neural substrates. More specifically, the question remains whether AD genetic risk factors exert their effects only at later life stages resulting in brain atrophy and clinical symptoms, or do they already impact neural substrates much earlier, i.e. through the entire lifespan and in the population at large? We hypothesized that a relation would be present throughout the lifespan, with higher AD-PGS, including presence of *APOE* ε4, showing association with lower hippocampal volumes early in life, as an offset effect. We tested whether AD genetic risk factors derived from genome-wide association studies (GWAS) [2] had an effect on hippocampal volume and volumetric changes in cognitively healthy individuals through the lifespan, and whether genetic risk interacted with age.

## 2. Methods

### 2.1. Sample

A total of 2690 valid scans from 1181 cognitively healthy participants, 4.1 to 95.7 years of age (mean visit age 39.7 years, SD 26.9 years), were drawn from five Norwegian studies. The studies included four sub-studies coordinated by the Center for Lifespan Changes in Brain and Cognition (LCBC); The Norwegian Mother and Child Cohort Neurocognitive Study (MoBa) [15], Neurocognitive Development (ND)[16], Cognition and Plasticity Through the Lifespan (CPLS) [17], Neurocognitive Plasticity (NCP) [18], and a study run collaboratively by LCBC and Oslo University Hospital, Novel Biomarkers (NBM) [19] (see Supplementary Material (SM), for details). The majority of participants were followed longitudinally, scan intervals ranging 0.2-11.0 years (mean = 2.9 years, SD =2.6 years). The sample is partly overlapping with [13, 20]. All were screened, and dementia, previous stroke with sequela, Parkinson’s disease, and other neurodegenerative diseases likely to affect cognition were exclusion criteria, with additional inclusion and exclusion criteria being applied per study (for details see SM). The participants in the sub-studies run by LCBC [15–18], constituting the majority (n = 1095), were cognitively healthy community dwellers, but complete absence of health problems was not required for inclusion. Participants with common health conditions, such as moderately elevated blood pressure and being on hypertensive treatment, were not excluded. They were recruited in part by newspaper and online ads, and in part through the population registry cohort study MoBa (see SM for details). The participants in the NBM study [19] (n = 86 at baseline) were recruited among patients scheduled for elective gynecological (genital prolapse), urological (benign prostate hyperplasia, prostate cancer, or bladder tumor/cancer) or orthopedic (knee or hip replacement) surgery in spinal anesthesia, turning 65 years or older in the year of inclusion. Informed consent was obtained for all participants; in writing from those 15 years and older, and from parents of participants below 15 years of age, and participants 12 years and older also gave oral consent. The studies were approved by the Regional Ethical Committee of South East Norway.

Additional criteria for being included in the present analyses were 1) having valid genetic data, 2) being of European ancestry as determined by genetic analyses (see below), 3) having at least one valid anatomical MRI scan with successful automatic hippocampal segmentation (see below). Sample descriptions for the total sample binned by timepoints are given in Table 1. Additional descriptions including distribution of sub-study samples per timepoint is given in Supplementary table 1 (see SM). To check for possible recruitment bias/age selectivity, the correlations of age and AD-PGS and *APOE* ε4 allelic variation (coded as both 0,1, or 2 ε4 alleles, as well as absence or presence of ε4 allele, 0 or 1) were calculated. For AD-PGS and age, there was a modest negative correlation, r = −.04, p = .0482, suggesting some age selectivity. No significant correlations or trends towards such were observed for *APOE* ε4 allelic variation, correlations with age being r = −.01, p = .4942 for *APOE* ε4 coded as no (0)/yes (1), and r = .00, p = .9356 for *APOE* coded as no ε4 allele (0)/ one ε4 allele (1) / two ε4 alleles (2). This suggests that there was no substantial sample selection age-bias with regard to genetic risk for AD. The proportion of *APOE* ε4 carriers in the present study appears largely in line with that which would be expected for the population in a Scandinavian country [21, 22].

**Table 1.**
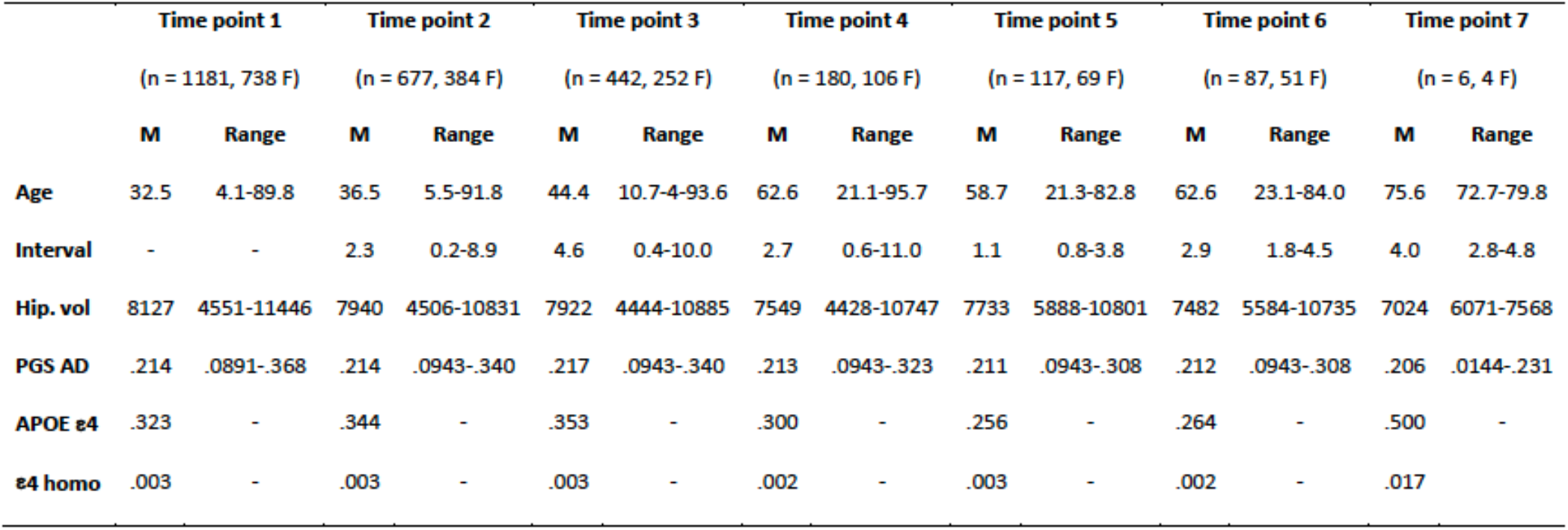
Sample descriptives. Age and interval are given in years. interval is interval since 1^st^ visit. Hippocampus volume (Hip. vol.) denotes number of voxels (mm^3^) in the hippocampal segmentation bilaterally. PGS AD = polygenic risk score for AD (threshold p <.5), given as a value 1^−03^, i.e. .300^−03^ is 0.000300. APOE ε4 denotes the proportion of participants with one or more Apolipoprotein E ε4 alleles. ε4 homozygotes (ε4 homo) denotes the proportion with two ε4 alleles.

|  | Time point 1 |  | Time point 2 |  | Time point 3 |  | Time point 4 |  | Time point 5 |  | Time point 6 |  | Time point 7 |  |
| --- | --- | --- | --- | --- | --- | --- | --- | --- | --- | --- | --- | --- | --- | --- |
|  | (n = 1181, 738 F) |  | (n = 677, 384 F) |  | (n = 442, 252 F) |  | (n = 180, 106 F) |  | (n = 117, 69 F) |  | (n = 87, 51 F) |  | (n = 6, 4 F) |  |
|  | M | Range | M | Range | M | Range | M | Range | M | Range | M | Range | M | Range |
| Age | 32.5 | 4.1-89.8 | 36.5 | 5.5-91.8 | 44.4 | 10.7-4-93.6 | 62.6 | 21.1-95.7 | 58.7 | 21.3-82.8 | 62.6 | 23.1-84.0 | 75.6 | 72.7-79.8 |
| Interval | - | - | 2.3 | 0.2-8.9 | 4.6 | 0.4-10.0 | 2.7 | 0.6-11.0 | 1.1 | 0.8-3.8 | 2.9 | 1.8-4.5 | 4.0 | 2.8-4.8 |
| Hip. vol | 8127 | 4551-11446 | 7940 | 4506-10831 | 7922 | 4444-10885 | 7549 | 4428-10747 | 7733 | 5888-10801 | 7482 | 5584-10735 | 7024 | 6071-7568 |
| PGS AD | .214 | .0891-.368 | .214 | .0943-.340 | .217 | .0943-.340 | .213 | .0943-.323 | .211 | .0943-.308 | .212 | .0943-.308 | .206 | .0144-.231 |
| APOE $\epsilon 4$ | .323 | - | .344 | - | .353 | - | .300 | - | .256 | - | .264 | - | .500 | - |
| $\epsilon 4$ homo | .003 | - | .003 | - | .003 | - | .002 | - | .003 | - | .002 | - | .017 | - |

### 2.2. Genotyping

Buccal swab and saliva samples were collected for DNA extraction followed by genome-wide genotyping using the “Global Screening Array” (Illumina, Inc). For a full description of genotyping and post-genotyping methods, including QC and imputation of untyped markers please see SM. AD-PGS of our sample was calculated using the allelic effect sizes from Lambert et al. [2]. The SNPs common to our data and Lambert et al. [2] were pruned to be nearly independent using the program PLINK 1.9 [23] with the following parameters: --clump-p1 1.0 ‒clump-p2 1.0 ‒clump-kb 500 ‒clump-r2 0.1. The linkage disequilibrium (LD) structure was based on the European subpopulation from the 1000 Genomes Project Phase3 [24]. Due to the complexity of the MHC region (build hg19; chr6:25,652,429-33,368,333), we removed SNPs in this region except the most significant one before pruning. Previous studies [25–27] have shown that PGS constructed using SNPs with association p value < 0.5 from Lambert et al. [2] have the largest effect on the risk of AD. Hence, we used the same threshold in the pruned set for computing the AD-PGS. Due to its known large effect, we computed AD-PGS with and without markers in the *APOE* region (build hg19; chr19:44,909,011-45,912,650). To test the effect of *APOE* itself we modelled the counts of *APOE* ε4 alleles directly by determining haplotypes of the two SNPs rs7412 and rs429358 [28, 29], coded as 0, 1, or 2 copies of the ε4 allele. We computed the genetic ancestry factors (GAFs) using principal components methods [30]. For the present analysis, only participants of European ancestry were included, excluding 89 persons for whom we had genotype data (see SM for further details). Finally, to investigate consistency of results across different p-value thresholds, analyses were recomputed limiting markers to those showing genome-wide significant association (i.e. p < .5e-08) with AD risk in Lambert et al. [2].

### 2.3. MRI data acquisition

Participants were scanned at a total of 4 Siemens scanners at 2 sites (1: Oslo University Hospital, 2: Curato (Currently Aleris), Oslo): A 1.5T Avanto equipped with a 12 channel head coil (Site 1 and 2), a 3T Skyra equipped with a 24-channel Siemens head coil (Site 1) or a 3T Prisma equipped with a 32 channel head coil (Site 1) (all Siemens Medical Systems, Erlangen, Germany). The pulse sequence used for morphometric analyses were one to two 3D sagittal T1-weighted MPRAGE sequences. For details on the pulse sequences used at each scanner, see SM. Other MRI volumes were recorded including sequences intended for and examined by a radiologist, to rule out and medically follow up incidental neuroradiological findings. Distribution of scans from the different scanners per timepoint is given in Supplementary table 1 (see SM).

### 2.4. Image analysis

All scans were reviewed for quality and automatically corrected for spatial distortion [31, 32]. Images were first automatically processed cross-sectionally for each time point with the FreeSurfer software package (version 6.0). This processing includes automatic hippocampal volumetric segmentation [33, 34]. In older subjects, FreeSurfer is shown to calculate consistent hippocampal volumes with reproducibility errors of 3.4%- 3.6% [35]. To extract reliable longitudinal subcortical volume estimates, the images were run through the longitudinal stream in FreeSurfer [36, 37]. Participants followed-up on different MRI scanners were independently processed for each scanner. To allow assessment of differences between scanners, 24 participants were scanned on all three scanners from Oslo University Hospital on the same day. Linear regression analyses were run testing the concordance between hippocampal volumes between scanners, yielding excellent agreement (Avanto vs Prisma R^2^ = .93; Prisma vs Skyra R^2^ = .94; Prisma vs. Avanto R^2^=.90). Thus, including scanner as covariate in the analyses would almost perfectly account for any possible scanner bias.

### 2.5. Statistical analyses

Analyses were run in R [38] version 3.6.0. General Additive Mixed Models (GAMM) using the package “mgcv” [39] version 1.8-28 were used to derive age-functions with a random intercept term per participant. Hippocampal volumes were predicted from 1) a smooth function of age and a linear function of AD-PGS, 2) a smooth function of age and linear functions of AD-PGS and presence/absence of one or more *APOE* ε4 alleles, 3) smooth functions of age and AD-PGS as well as a tensor interaction term of age and AD-PGS, 4) a linear function of presence/absence of one or more *APOE* ε4 alleles and a smooth interaction between age and presence/absence of one or more *APOE* ε4 alleles. Marginal maximum likelihood was used for smoothness selection. In all models, scanner, intracranial volume, sex and the first 5 GAFs, in addition to genotyping batch (1; n = 1014, 2; n = 166) were entered as covariates. See SM for further details.

## 3. Results

In the lifespan sample, hippocampal volumes increased in early development and declined in older age, as shown in Figure 1A. We found a significant negative effect of AD-PGS on hippocampal volume (t = −2.017, p = .0437), as shown in Figure 1B. There was no significant age interaction with AD-PGS (F = 1.487, p = .1915). While the analysis was based on the continuous AD-PGS score, inspection of the age trajectories for the first and fourth quartile values of AD-PGS (Figure 1C), did not give any consistent indication of greater effects of higher PGS for AD with age. Rather, while there was time after age 60 when the hippocampal curves merge for the different subgroups, they were otherwise mostly ordered throughout life so that those with the lowest quartile PGS for AD had the highest volume, while those with the highest quartile PGS for AD have the lowest volume. Of note, the absolute volume difference between those with lower and higher AD-PGS appeared to be about the same in young adults (∼20s) as in the oldest adults (∼80s).

**Figure 1.**
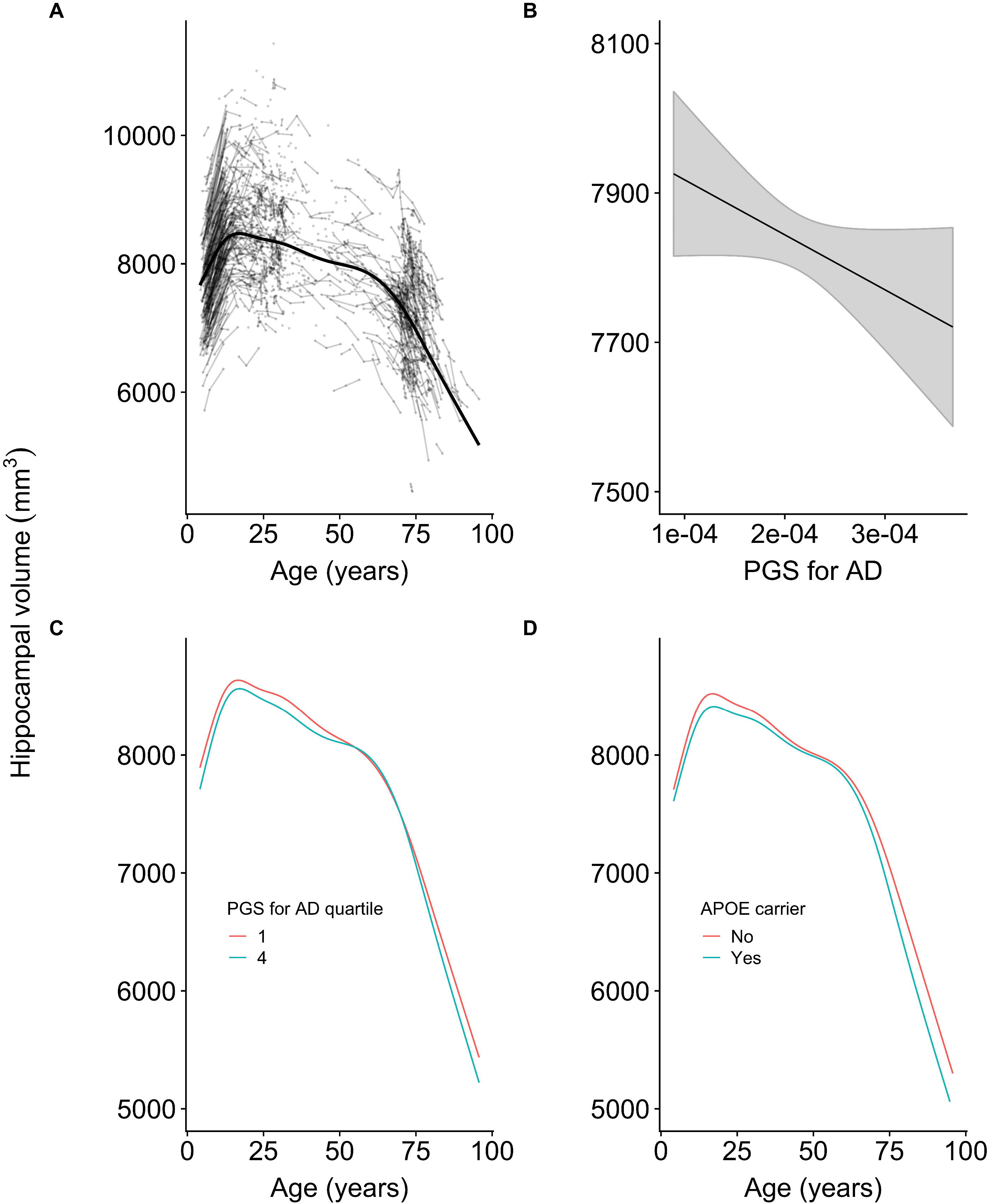
Hippocampal volume (across hemispheres, shown in mm^3^ on the Y-axis) and change in relation to A) age (in years, x, axis) plotted with individual trajectories overlaid, B) polygenic score (PGS) for AD (x-axis, continuous scale 0-1), C) age (in years, x axis) with PGS for AD set to the first (red line) and fourth quartile (blue line) of the sample, and D) trajectories for carriers (blue line) and non-carriers (red line) of the APOE ε4 allele.

Analysis with *APOE* status as the predictor of hippocampal volume likewise showed a significant negative effect of presence of the ε4 allele (t = −2.779, p = .0054). There was no significant age interaction for *APOE* allelic variance, but a trend was observed (F = 2.103, p =.075). This is in line with ε4 allele carriers having smaller hippocampal volumes in young age, as an offset effect. Importantly, the trend towards age interaction cannot be interpreted as indicating faster atrophy in older age, as carriers in young adulthood (∼20s) appeared to display lower hippocampal volumes similar to carriers in older adulthood (∼80s; Figure 1D).

AD-PGS excluding the *APOE*-region did not show a significant effect on hippocampal volume, although a trend in the same direction as the full model was observed (t = −1.744, p = .0812). There was no evidence for an age interaction upon utilizing the AD-PGS excluding markers in the *APOE* region (F = 0.985, p = .4670).

Recomputing the analyses with an AD-PGS limited to SNPs only showing genome-wide significant association (i.e. p < 5e-08) with AD risk in Lambert et al. [2], confirmed a significant negative effect of AD-PGS on hippocampal volume (t = −2.444, p = .0146). This result was reduced to a trend when excluding markers in the *APOE* region (t = −1.699, p = .0893). However, significant age interactions appeared in the PGS analyses limited to genome-wide significant markers, both with and without markers in the *APOE* region (with *APOE*: F = 2.269, p = .0335; without *APOE*: F=2.211, p = .0147; Supplementary Figures 1 and 2, respectively). Especially when excluding the *APOE* region, there seemed to be a somewhat more negative effect of higher AD-PGS on hippocampal volume in older age (above 80 yrs).

## 4. Discussion

This study shows that genetic risk for AD is associated with effects on hippocampal volume throughout life, having neurodevelopmental offset effects observable from childhood in cognitively healthy well-functioning participants. Higher AD-PGS and carrying the *APOE* ε4 allele specifically, were associated with lower developmental hippocampal volume offsets, and persons with higher genetic risk for AD remained having lower hippocampal volumes with age. Notably, this main effect was consistently observed for PGSs constructed using SNPs with two different association p values [2].The association of hippocampal volume and AD-PGS through the lifespan is a novel finding, as to our knowledge no previous reports exist on the lifespan trajectories of polygenic AD risk at an endophenotypic level. However, and as expected, the effect was not large, and when using the AD-PGSs computed without the *APOE* region as predictor, the effect was reduced to a trend.

Having low versus high genetic risk for AD was associated with roughly equivalent difference in hippocampal volume at the age of 25 and 80 years. The effect of the PGS constructed using SNPs with association p value < 0.5 from Lambert et al. [2], shown to have the largest effect on the risk of AD [25–27], did not significantly interact with age. However, the PGS effect was not equally apparent at all ages among older adults, and notably, age interactions were observed for the same PGS constructed using SNPs with the lowest association p-value. Possibly, different extent of effects at different ages may illuminate why one small recent study found associations of an AD genetic risk score and hippocampal atrophy over a two-year interval in healthy older adults (n = 45), though no association with individual variation in volume was seen at baseline (n = 66) [7]. AD-PGS did correlate modestly with age in our sample, and there may be some sample-selection bias in healthy older age samples, so it would be interesting to see whether age interactions could be consistently observed if persons were followed for even longer and at higher ages. We cannot, based on the current sample, exclude a mixture of neurodevelopmental offset and aging effects, though the neurodevelopmental offset effects were the most consistently supported by the present results.

There was only a trend for effects of *APOE* allelic variants to interact with age, and this trend was not clearly indicative of faster atrophy for *APOE* ε4 carriers in older age. Cross-sectional studies have shown that *APOE* ε4 [3, 4] carriers tend to have lower hippocampal/medial temporal lobe volumes at various ages also in healthy persons [5], and notably, this has been identified even in neonates [40]. However, it has been difficult to interpret whether these developmental structural brain differences actually do represent long term risk factors, as longitudinal imaging data have been lacking. The only such study we know of was restricted to older adults aged 55-75 [6], showing greater hippocampal atrophy in ε4 carriers across a five year interval. The present study hence partly confirms, yet nuances and extends previous reports, in showing a main effect through the lifespan, i.e. also an offset effect.

The present study adds to the evidence for early life factors exerting a continuous influence on later-life function [11, 12], showing that genetic factors established to influence late life neural and cognitive disease work in part in a temporally stable and dimensional manner. That is, genetic risk factors for AD seem not to only manifest at late life in neural atrophy and clinical symptoms, but appear to start influencing neural substrates of cognition early on, through the entire lifespan and in the population at large. In this regard, currently observed effects of the *APOE* ε4 allele cannot well be interpreted only within the framework of antagonistic pleiotropy [8–10], as this allelic variant appears to have in part similar effects on neural substrates of memory function and different ages. As for AD-PGS, currently observed effects are of similar magnitude in young adulthood as in much of older adulthood, meaning that explanations evoking different effects at neural substrates at different ages may be incomplete.

This does not necessarily imply that effects on behavior may be readily observed through the lifespan. A number of studies have indeed not observed effects of *APOE* status on standard neuropsychological memory tests [41, 42]. However, as recently reported, this does not mean that effects of genetic risk for AD do not manifest early, in more fine-grained behaviors dependent on hippocampal circuitry, such as spatial navigation [42]. A related, yet different account, is the magnification or resource modulation hypothesis [43–45], stating that genetic effects are magnified in persons with constrained neural resources, such as older – and putatively also developing – individuals. Here, it is assumed that the function relating brain resources to cognition is non-linear, so that genetic differences exert increasingly larger effects on performance with lesser neural substrates. Regardless of direct effects on behavior, smaller hippocampi can within these various accounts be seen as a risk factor. Within a brain reserve account, such differences in brain structure may relate to differences in tolerance to pathology before one falls under a functional threshold [46]. Hence, these differences may become readily functionally apparent only in older age, but they likely are there to begin with and throughout life in a stable manner. This supports a lifespan model of dementing disorder [47], where effects of common genetic variants in part work in a stable manner to be one of several factors affecting risk for cognitive decline and neurodegenerative disease.

As risk is multifactorial [47, 48], the genetic component is one of many factors that will be at play. It may be especially important to target other risk factors through life for those who are, albeit well-functioning, at the highest genetic risk. Such other risk factors for which interventions should be offered may, according to the new World Health Organization (WHO) guidelines for Risk Reduction of Cognitive Decline and Dementia [48], include physical activity level, tobacco use, diet, hypertension and diabetes management [48]. While the recommendations are partly based on data from experimental interventions, much literature on risk factors is based on observational data. We do not know if such factors are really protective. It may be that the risk and protective factors are markers of some other favorable, and perhaps genetic, trait [49, 50]. Still, we note that WHO calls for further research in at-risk populations, and research to understand how timing affects the impact of interventions on cognitive decline and dementia. Based on the present results for genetic risk and hippocampal volume and its age change, risk is not something that only has an impact at a specific age, such as midlife or late life, but really may work in part in a continuous manner through all of life. If correct, a possible implication from these data is that attempts to reduce risk for neurodegenerative disease should be aimed at the entire lifespan.

In conclusion, endophenotypic expression of genetic risk for AD may be seen in a dimensional and lifespan perspective, not being confined to clinical populations or older age. This emphasizes that a broader population and age range are relevant targets for attempts to prevent AD. Future studies should investigate which genetic factors established to influence AD work in a temporally stable manner, across manifestations of health and disease, and which may have more pronounced effects in aging, to help define targets for prevention.

## Acknowledgements/Conflicts/Funding Sources

We are grateful to all participants for their time and commitment, and to all colleagues in LCBC for taking part in the gathering and preprocessing of the data. We thank Mrs. Tanja Wesse, Mrs. Sanaz Sedghpour Sabet, and Dr. Michael Wittig at the Institute of Clinical Molecular Biology, Christian-Albrechts-University of Kiel, Kiel, Germany for technical assistance with the GSA genotyping. The LIGA team acknowledges computational support from the OMICS compute cluster at the University of Lübeck. This research was funded by grants from the Norwegian Research Council (to KBW, AMF and YW), the National Association for Public Health’s dementia research program, Norway (to A.M.F.), and the European Research Council’s Starting (grant agreements 283634, to A.M.F. and 313440 to K.B.W.) and consolidator Grant Scheme (grant agreement 771355 to KBW and 725025 to AMF). The funding sources had no role in the study design; in the collection, analysis, and interpretation of the data; in the writing of the report; or in the decision to submit the article for publication.

