## Supplementary_material for "Genetic risk for Alzheimer’s disease predicts hippocampal volume through the lifespan"

**Methods**

**Sample**

The studies from which the present sample was drawn used slightly different inclusion and exclusion criteria, as they had different aims and objectives. The Norwegian Mother and Child Cohort Neurocognitive Study (MoBa) [1] recruited children through the population registry cohort study MoBa, at observation mean age was 8.6 years (SD = 2.9 yrs), and age range was 4.1-16.2 years. Neurocognitive Development (ND)[2], Cognition and Plasticity Through the Lifespan (CPLS) [3] recruited through newspaper and online ads, and Novel Biomarkers (NBM) [4] recruited from a non-neurological disease patient population as described. MoBa, ND, CPLS and NMB were merely observational studies, whereas the NCP [5] study included one or more 10 week periods of memory training or rest, with scans in between these. All persons in the NCP project were offered memory training, irrespective of genetics, and hence, this should not constitute a bias. All participants were compensated a modest sum for their participation, depending on amount of examinations (for the structural scan session around NOK 500, or USD $60). Education was recorded as number of years of education to the highest attained degree for adults (age >= 18 years), and for participants below 18 years of age, the average of paternal and maternal years of education to the highest attained degree was entered, or if unavailable, for one parent (either available). By this measure, education was obtained in a comparable manner for most participants (n = 1000, mean = 16.3 years, SD = 2.8 years, range 7-24 years).

While the proportion of $\varepsilon$4 allele carriers, as given in Table 1, was somewhat higher at time points 1-4 than at time points 5-6, as seen in Table 1, this should not be interpreted as selective drop-out of $\varepsilon$4 carriers. Only participants of the NCP project were offered more than four scans, and they also constitute the majority of participants at time point 4 (see SM). The proportion of $\varepsilon$4 carriers in the NCP sample was lower than that of the combined sample at baseline, namely .274, and so remained relatively similar across timepoints.

**Supplementary Table 1**

Distribution of number of participants from different sub-studies and scanners across time points (Tp).

| | Tp 1 | Tp 2 | Tp 3 | Tp 4 | Tp5 | Tp6 | Tp7 |
|---|---|---|---|---|---|---|---|
| **Sub-study** | | | | | | | |
| MoBa | 223 | 195 | 105 | - | - | - | |
| NBM | 86 | 81 | 62 | 27 | - | - | |
| ND | 224 | 130 | 63 | - | - | - | |
| CPLS | 462 | 102 | 67 | 22 | - | - | |
| NCP | 186 | 169 | 145 | 131 | 117 | 87 | 6 |
| **Scanner** | | | | | | | |
| Avanto 1 | 416 | 352 | 123 | - | - | - | - |
| Avanto 2 | 86 | 81 | 62 | 27 | - | - | - |
| Prisma | 100 | 37 | 105 | - | - | - | - |
| Skyra | 579 | 207 | 152 | 153 | 117 | 87 | 6 |

**DNA handling, genotyping, and data processing**

*DNA extraction, QC and genotyping*. Prior to genotyping, DNA was extracted either in Oslo from saliva or buccal swabs (n=592), or at the LIGA laboratory at University of Lübeck for saliva samples (n=416, incl. 2 technical duplicates) and buccal swabs (n=586, incl. 1 technical duplicate). All DNA samples were then quantified, quality controlled, normalized, and aliquoted (to approx. 35ul at ~50ng/ul) in Lübeck leading to a total of 1,594 DNA samples (incl. 3 technical duplicates) that were subjected to genotyping using the Global Screening Array (GSA; Illumina, Inc.) with shared custom content. Genotyping was performed in two batches (batch 1: n = 1,402, batch 2: n = 192) at the Institute of Clinical and Molecular Biology at UKSH Campus Kiel on an iScan instrument following the manufacturer's instructions. Three samples from batch 1 (2 buccal swabs, 1 saliva) failed genotyping leaving n=1,591 for further processing.

*Post-genotyping data processing, QC and imputation*. All data processing steps were performed in the LIGA laboratory in Lübeck. Genotype calling was performed in GenomeStudio v2.0.4 using manifest "GSAsharedCUSTOM_20018389_A6" (Illumina, Inc.) providing annotations for a total of 696,375 variants. GenTrain v3.0 (Illumina, Inc.) was used for automatic clustering and genotype calling. Quality assessments at this stage revealed one sample failing QC from batch 2 (using call rate < .95 and p50 GC < .7 as threshold) so that all 1,590 samples were exported using the PLINK Input Report 2.1.4 module of GenomeStudio. Subsequent data processing used an automated workflow developed in LIGA executed in the high-performance computing environment ("OmicsCluster") available at University of Lübeck. This entailed exclusion of 110,579 variants with GenTrain values <.70 [6] in a reference dataset of ~20,000 DNA samples genotyped in a separate project, conversion of variant alleles to forward (plus) stand using PLINK (v1.90b4;[7]; command: '--flip') and checking for inconsistencies between reported and genetic sex ('--check-sex'). At this stage, 13 samples (batch 1: n=11, batch 2: n=3) were excluded because genetic sex could not be determined unambiguously from the genotype data, and non-matching sex information was recorded for re-inspection of clinical data for 29 samples (batch 1: n = 25, batch 2: n=4). Subsequent sample- and variant-level QC entailed filtering with PLINK commands '--mind 0.05 --geno 0.02 --hwe 0.000005 --maf 0.01', resulting in 446,837 high-quality variants with MAF ≥1% across 1,562 (batch 1: n=1,381, batch 2: n=181) samples. These data were used to create an LD pruned dataset with '--indep-pairwise 1500 150 0.2 --maf 0.05' followed by pairwise genetic similarity analyses using '--Z-genome --min 0.06' to identify cryptic relatedness. All three technical replicates were identified correctly and showed PI_HAT values of 1. In addition, samples with >3 standard deviations of pairwise matches at PI_HAT>0.06 were excluded (batch 1: n=2 samples, batch 2: n=0). The LD pruned dataset was then used for principal component analysis (PCA; using PLINK command '--pca') along with the reference dataset of the 1000 Genomes Project Phase 3 (1000G,[8]) to assign ethnic descent groups using the five 1000G super- populations by k-nearest neighbor (k-NN; k=9) classification (using R package 'class' in R 2.3.2; [9]). Subsequently, data were recoded into VCF format ('--recode vcf-iid'), whereby ambiguous SNPs were removed, and strand mismatches re-checked and corrected with BCFtools 1.9 ( '+fixref -m flip -d -f GRCh37.fasta') and confirmed ('+af-dist'). SHAPEIT2 (v2.r837; [10]) was used for phasing. Variants not matching to the reference haplotypes were excluded ('-check -M), all remaining variants were phased with the same reference data, i.e. the genetic map of the 1000G and the HRC reference panel haplotypes Release 1.1 (EGAD00001002729). The phased genotype data were then subjected to imputation using the HRC reference with Minimac3 [11] applying default parameters. Overall, this procedure resulted in 39,131,578 genotypes across 1,560 (batch 1: n=1,379, batch 2: n=181) (incl. all 3 technical replicates) samples.

*Polygenic score computation.* Imputed dosages were further quality controlled by: 1.) removing non-European subjects determined by the PCA analyses; 2.) SNPs having imputation R square <0.8; and, 3.) minor allele frequencies (MAF) <0.05. The resultant dosage genotypes were converted to best-guess genotypes, i.e. 0, 1, or 2 copies of the minor allele for each SNP. In total, 5.2 million SNPs remain. AD GWAS summary statistics were downloaded from http://web.pasteur-lille.fr/en/recherche/u744/igap/igap_download.php with permission. Shared SNPs between our best-guess genotype dataset and the GWAS summary statistics were pruned to be near independent with PLINK using parameters --clump-p1 1.0 --clump-p2 1.0 --clump-kb 500 --clump-r2 0.1 and LD structure from 1000G. To avoid the impact of the complex LD structure of the MHC region (build hg19; chr6:25652429-33,368,333), only the most significant SNPs of this region were included. We computed PGS with and without the APOE region to investigate its effect on the hippocampus. SNPs with $p < 0.5$ and $p < 5e-08$ in the pruned set were used for constructing the PGS for our samples.

*Genetic ancestral factors (GAF)*. The pre-imputation QC'ed genotypes were used for estimating GAF for European samples (as determined by the k-NN approach on 1000G superpopulations, see above). SNPs with MAF < 0.1 were excluded first and then pruned to be nearly independent by PLINK using parameters, --indep-pairwise 100 50 0.1. Then, the remaining SNPs were included in the GAF estimation with the PLINK command, --pca. The top 20 principal components were retained to be used as co-variates in the statistical analyses.

**MRI data acquisition**

The pulse sequence used for morphometric analyses were one to two 3D sagittal T1-weighted MPRAGE sequences. Avanto site 1 and 2: 160 slices, repetition time (TR), 2400ms; echo time (TE), 3.61ms/3.79ms (Site 1/2); time to inversion, 1000ms; flip angle, 8°; matrix, 192x192; field of view, 240; voxel size, 1.25x1.25x1.20 mm per participant per visit. Scanning time for each MPRAGE sequence was 7min 42s. Skyra: 176 slices, TR = 2300 ms, TE = 2.98 ms, flip angle = 8°, voxel size = 1 × 1 × 1 mm, FOV= 256 × 256 mm. Prisma: 208 slices, TR = 2400 ms, TE = 2.22 ms, TI = 1000 ms, flip angle = 8°, voxel size = 0.8 x 0.8 x 0.8 $mm^3$, FOV = 240 x 256 $mm^2$.

**Image Analysis**

All scans were reviewed for quality and automatically corrected for spatial distortion due to gradient nonlinearity [12] and B1 field inhomogeneity [13]. Images were first automatically processed cross-sectionally for each time point with the FreeSurfer software package (version 6.0; http://surfer.nmr.mgh.harvard.edu/). This processing includes motion correction, removal of non-brain tissue, automated Talairach transformation, intensity correction and volumetric segmentation [14]. The segmentation procedure automatically labels each voxel in the brain as one of 40 structures [14], using a probabilistic brain atlas specific for the current image acquisition protocol [15]. In older subjects scanned on the same Siemens MRI scanner, FreeSurfer is shown to calculate consistent hippocampal formation volumes from 1.5T MPRAGE images with reproducibility errors of 3.6% and 3.4% for the left and right hippocampus respectively [16]. To extract reliable longitudinal subcortical volume estimates, the images were run through the longitudinal stream in FreeSurfer [17]. Specifically, an unbiased within-subject template volume based on the cross-sectional images was created for each participant, and processing of all time points was then initialized using common information from this template. This increased sensitivity and robustness of the longitudinal analysis and ensured inverse consistency [18]. In addition, new probabilistic methods (temporal fusion) were applied to further reduce the variability across time points. Participants followed-up on different MRI scanners were independently processed for each scanner. To allow assessment of differences between scanners, 24 participants were scanned on all three scanners from Oslo University Hospital on the same day. Linear regression analyses were run testing the concordance between hippocampal volumes between scanners, yielding excellent agreement (Avanto vs Prisma $R^2$ = .93; Prisma vs Skyra $R^2$ = .94; Prisma vs. Avanto $R^2$=.90). Thus, including scanner as covariate in the analyses would almost perfectly account for any possible scanner bias.

**Supplementary results**

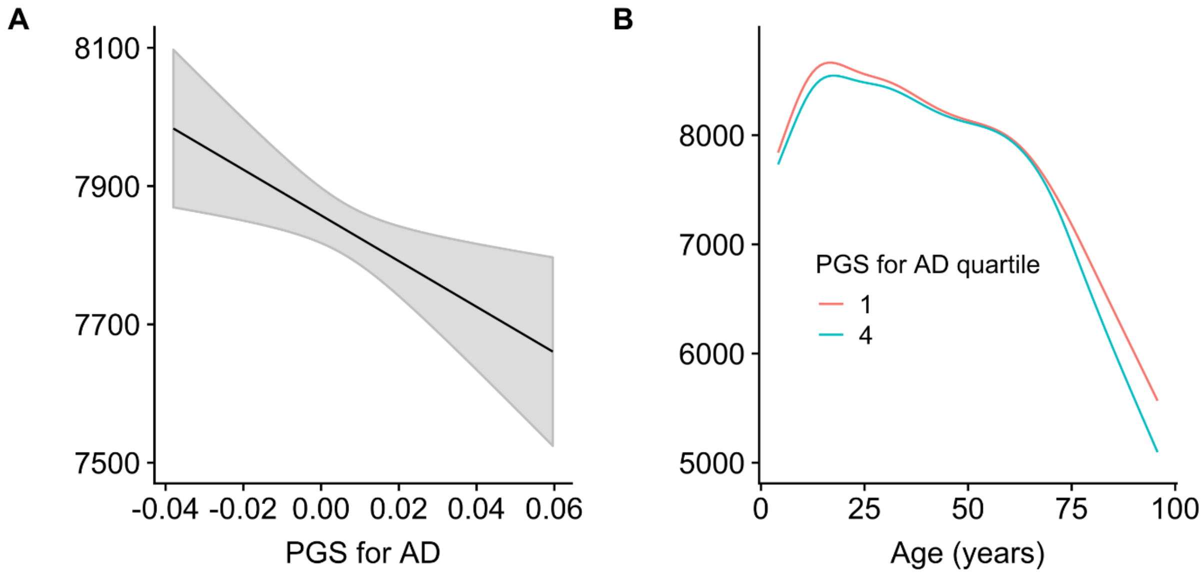


**Supplementary figure 1.** Hippocampal volume (across hemispheres, shown in $mm^3$ on the Y-axis) and change in relation to A) AD-PGS including the APOE region as calculated based on effect sizes from Lambert et al. [19] at p < . 5e-08 (x-axis, continuous scale 0-1). B) Age (in years, x axis) with AD-PGS at p < .5e-08 including the APOE region set to the first and fourth quartile of the sample.

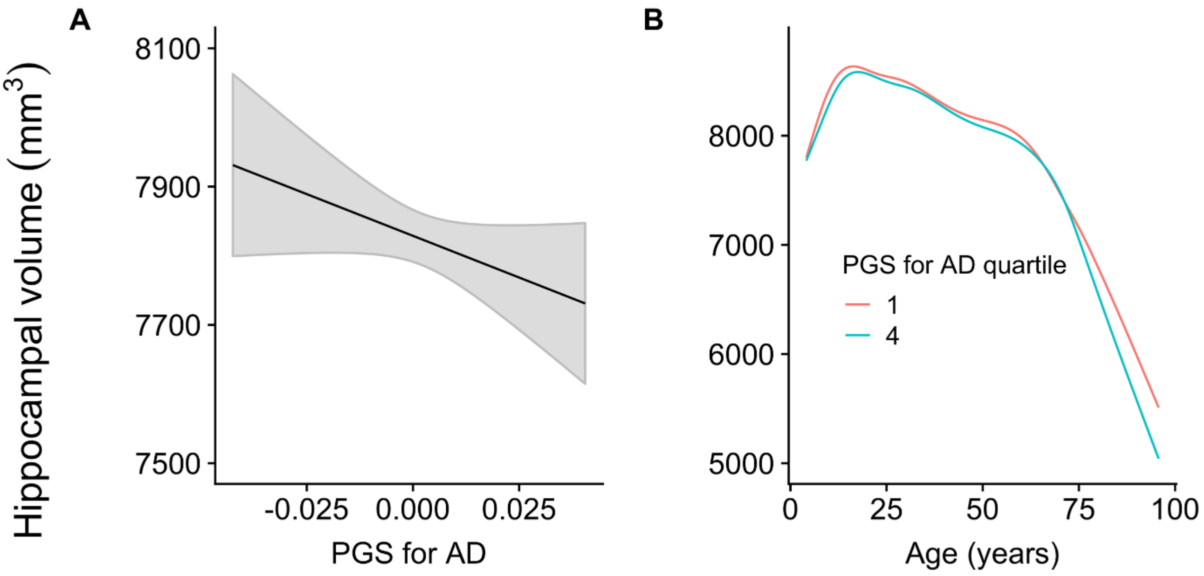


**Supplementary figure 2.** Hippocampal volume (across hemispheres, shown in $mm^3$ on the Y-axis) and change in relation to A) AD-PGS excluding the APOE region as calculated based on effect sizes from Lambert et al. [19] at p < . 5e-08 (x-axis, continuous scale 0-1). B) Age (in years, x axis) with AD-PGS at p < . 5e-08 excluding the APOE region set to the first and fourth quartile of the sample.
